# Development of a recombinant adeno-associated virus vector for human T lymphocyte- and natural killer cell-targeted gene therapy

**DOI:** 10.64898/2026.02.26.707014

**Authors:** Hendrik Jahnz, Martin V. Hamann, Hongil Kim, Yaping Sun, Natalia Salazar Quiroz, Li Zhu, Umm E Swaiba, Daniel Foth, Niklas Beschorner, Priti Kumar, Ulrike C. Lange

## Abstract

Recombinant adeno-associated virus (rAAV) vectors are widely used for gene delivery but show limited efficiency in immune cells, including T lymphocytes and natural killer (NK) cells. To overcome this barrier, we developed a modular rAAV vector engineering strategy that integrates capsid retargeting with genome optimization. We report a CD7-targeted rAAV vector (CD7-AAV6/9) featuring a nanobody-fused hybrid capsid derived from a rationally selected chimeric combination of AAV6 and AAV9. CD7-AAV6/9 enables efficient and selective transduction of immortalized and primary human T and NK cells *in vitro* and *in vivo* in a humanized mouse model, achieves high production titers, and exhibits markedly reduced off-target transduction compared with wild-type serotypes. In parallel, we demonstrate that incorporation of a human gene–derived intron into the vector genome overcomes host-mediated transcriptional repression and enables robust transgene expression in human CD7⁺ T lymphocyte and NK cell populations. To our knowledge, this represents the first application of intron-mediated enhancement in a rAAV vector context. Together, our findings establish an integrated capsid–genome design framework for targeting human T and NK cells, notoriously challenging immune cell populations for gene therapy, and provide a versatile platform readily adaptable to alternative surface markers and therapeutic payloads.

## 1. Introduction

Adeno–associated viruses (AAVs) have emerged as one of the leading platforms for *in vivo* gene therapy, with multiple recombinant AAV (rAAV) products now approved for clinical use, for instance, in inherited retinal disease, spinal muscular atrophy and hemophilia [1–5]. rAAV vectors are considered safe for human use as wild–type AAVs are not associated with human diseases. AAVs are replication–incompetent, and the vector genome predominantly persists as episomal DNA. In addition, AAVs can transduce both dividing and non–dividing cells, support long–term transgene expression and are highly amenable to capsid and genome engineering [6–12].

Naturally occurring AAV serotypes display broad tropism, leading to efficient infection of cells in organs such as liver, kidney, muscle and central nervous system [4,10,13,14]. This broad tropism can however cause substantial off–target exposure necessitating high doses when used in recombinant vector format, and has been implicated in dose–limiting toxicities and adverse events in clinical trials [6,15]. A major translational bottleneck in the rAAV field is achieving efficient and selective cell/tissue-specific delivery while minimizing transduction of non–target tissues *in vivo* [16,17]. Furthermore, most naturally occurring AAV serotypes show poor transduction of lymphocytes, with AAV6 being the most effective, yet non-specific, serotype, as it also targets epithelial and muscle cells and most noteworthily, hepatocytes. Given the rapid advances in the field of immune therapy (e.g. immune check point therapy, CAR T/NK), the demand for gene therapy vectors targeting immune cells, particularly T and NK cells for CAR T cell therapy, is expected to increase considerably in the near future.

AAV capsid engineering to alter vector tropism has emerged as powerful strategy to address limitations linked to non-specific tropism. Approaches range from high–throughput selection of diversified AAV libraries to rational design approaches based on known receptor–ligand interactions [10,18–23]. One rational strategy is the display of targeting ligands on exposed AAV capsid surface loops, for example, nanobodies (Nbs) which are heavy–chain–only antibody fragments that bind surface receptors specific to designated target cells [16,18,24]. Nanobody-engineered capsids can redirect rAAV vectors enhancing both efficacy and safety of *in vivo* cell gene therapy approaches [16,22].

We present here a rationally designed rAAV vector, termed CD7-AAV6/9-int, that combines capsid engineering for targeted tropism and transgene-cassette engineering for optimized expression in human T lymphocytes and NK cells. Specifically, in addition to an in-frame fusion of the AAV capsid to a CD7-specific nanobody, which enables targeting to human T and NK cells, the vector carries an intron-optimized transgene cassette to counteract host-mediated silencing of transgene expression. CD7-AAV6/9-int transduces CD7-positive CD4+/CD8+ T lymphocytes and NK cells *in vitro* and *in vivo* in humanized mice with markedly reduced off-target activity. We propose that the integrated ‘capsid–plus–genome’ design exemplified here provides a blueprint for next–generation *in vivo* gene therapy vectors for immune cell-based interventions.

## 2. Results

### 2.1. Engineering of AAV6/9 hybrid capsids with a nanobody targeted CD7 surface epitope

We previously developed nanobody capsid engineering as a powerful technique to redirect AAV target cell tropism towards CD4+ T lymphocytes [16]. Here we present a next generation CD7^+^-immune cell-targeted rAAV vector generated through a rational design strategy. First, we leveraged AAV capsid proteins from serotypes AAV6 and AAV9, selected for their complementary properties, efficiency for transduction of human lymphocytes (AAV6) and favorable systemic biodistribution profile and extensive clinical safety data (AAV9), and combined these features into a single hybrid capsid comprising capsid protein VP1 derived from AAV6 and capsid proteins VP2 and VP3 from AAV9 (Fig 1A). Next, we incorporated a nanobody targeting the human CD7 surface glycoprotein into the GH2/GH3 loop of VP1 (position T456) (Fig 1A). Incorporating the CD7-specific nanobody, in contrast to a CD4 nanobody, broadens tropism towards a range of lymphocyte subsets, including CD4⁺ and CD8⁺ T cells and NK cells [34]. Capsid-engineering yields consistently higher titers compared to most wild-type serotypes. Compared to wild-type AAV6, often considered vector of choice for lymphocyte transduction, we obtained ∼17 to 20 fold higher production titers for the CD7-nanobody capsid engineered AAV6 and AAV6/9 vectors respectively (Fig 1B). Structural modeling using AlphaFold predicted that CD7 nanobody incorporation did not alter VP capsid protein conformation (Fig 1C) and electron microscopy (EM) combined with immunogold-staining for nanobody showed natural capsid morphology for CD7-AAV6/9 particles and surface presentation of the CD7-Nb (Fig 1D).

**Figure 1:**
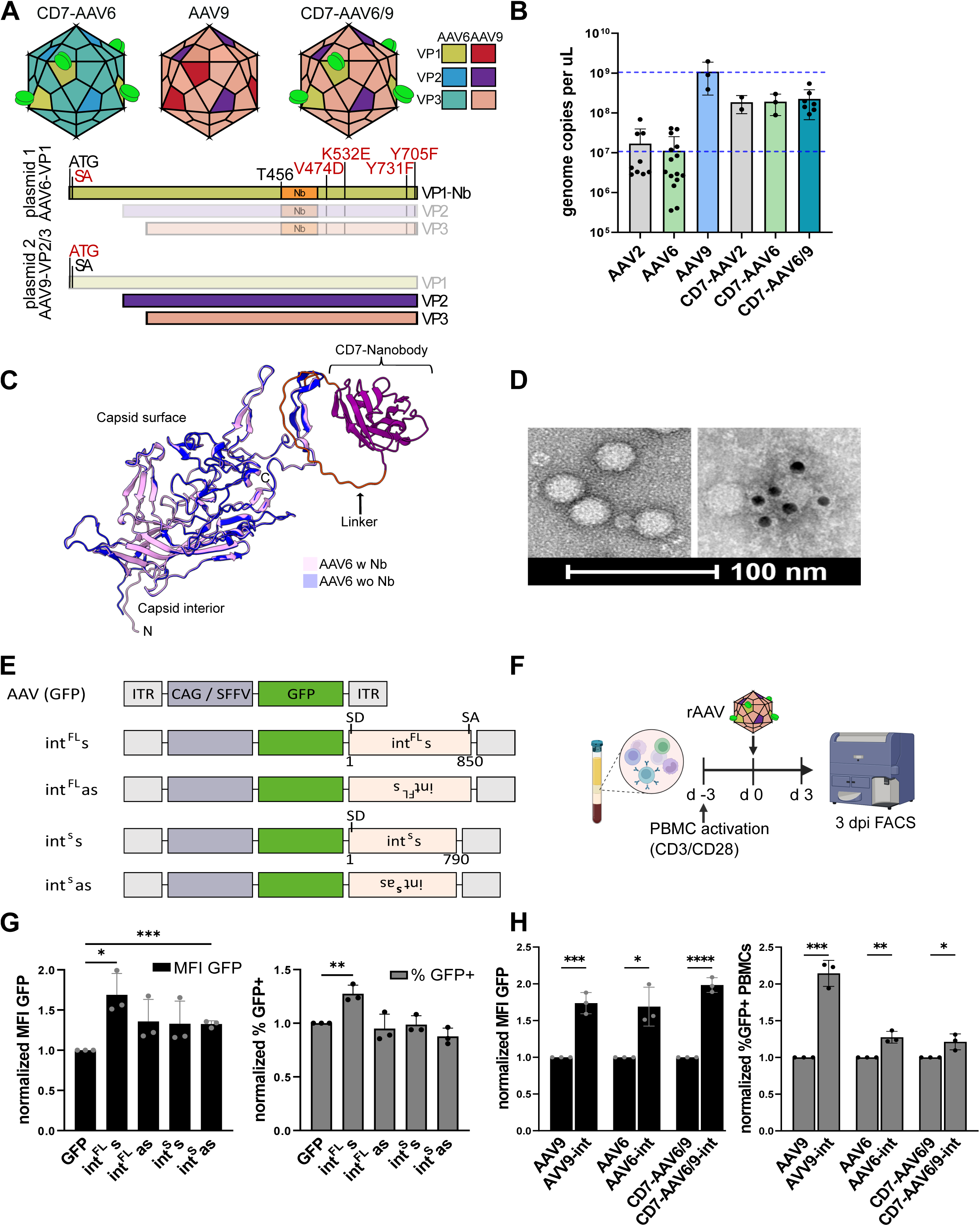
Generation and characterization of intron-optimized CD7-AAV6/9. **A)** Schematic representation of AAV capsids displaying CD7-Nanobody (Nb) (bright green) and VP1-3-coding constructs used for CD7-AAV6/9 production. The amino acid substitutions introduced are indicated in red; SA splice acceptor. **B)** Production titers of the different AAV serotypes. Shown are means with SD (n≥2). **C)** AlphaFold prediction superimposing AAV6 VP3 with (rose) and without (violet) CD7-Nb insertion. CD7-Nb is depicted in purple and GS-linker in orange. **D)** Representative negative-stained electron micrographs of CD7-AAV6/9 particles (left) and immuno-gold staining of the same particles for the Nb component (right). Scale bar is depicted. **E)** Schematic representation of rAAV transgene cassettes with intron (int) insertions. Intron length is indicated as base pairs. ITR (inverted terminal repeat), ^FL^ (full length), ^s^ shortened, s – sense, as (antisense), SD (splice donor), SA (splice acceptor). **F)** Experimental outline of rAAV PBMC transduction (10,000 gc/cell) and flow cytometry analysis at 3 dpi. **G)** Shown are normalized mean fluorescence intensity (MFI) (left) and % GFP-positive PBMCs (right) at 3 dpi relative to transgene without intron (mean ± SD, n=3). **H)** Same experimental setup as in C), but using capsid serotypes 9, 6 and hybrid 6/9, as indicated. All intron-containing vectors contain the same sense-full-length intron (int^FL^s). GFP MFI and % GFP-positive cells at 3 dpi normalized to transgene without intron. Shown are mean ± SD, n=3. For all panels, statistical analysis utilized the unpaired t-test, p < 0.05 (*), p < 0.01 (**), p < 0.001 (***), and p < 0.0001 (****).

### 2.2. Intron insertion optimizes AAV transgene cassette for expression and transduction rates in primary immune cells

Transcription of intronless transgenes, such as those delivered by rAAVs, is known to be subject to repression by the Human Silencing Hub (HUSH) complex, a key epigenetic regulator of exogenous DNA [43,44]. We reasoned that insertion of intronic sequences could shield rAAV transgenes from HUSH-mediated repression. To test this, we engineered four variants of a single-stranded AAV transgene encoding the GFP reporter gene. Each construct contained a variation of the second intron of the human β-globin gene (*HBB*) inserted downstream of the GFP coding sequence. The variants included either the full-length (850nt) intron (FL) or a shortened version (S, 790nt; 3’ truncated by 60 nucleotides and lacking the splice branch-point site [44]), each cloned in either sense (s) or antisense (as) orientation within the transgene (Fig 1E). The short intron does not support splicing and thus allows to distinguish between effects mediated by active splicing or due to intron presence alone. PBMCs were transduced with equal amounts of rAAV6 engineered with the above-described transgene variants and GFP expression determined by flow cytometry three days later (Fig 1F). All intron-containing constructs produced a marked increase in GFP mean fluorescence intensity (MFI) relative to intron-less controls (Fig 1G left panel). The full-length intron in the sense orientation (FL-s) additionally led to a significant increase in the percentage of GFP⁺ cells (Fig 1G, right panel). Taken together, transgene insertion of the full-length intron in sense orientation (FL-s) significantly improved expression performance of rAAV6 vectors.

To assess the generality of this effect to other AAV serotypes, we compared expression of AAV9, AAV6 and AAV6/9 capsid-delivered transgenes without or with FL-s intron insertion (AAV-int). Intron insertion led to enhanced transgene expression in all cases, suggesting that this effect is determined by host-cell regulatory mechanisms rather than by specific AAV capsid features (Fig 1H). Our data shows that intron insertion boosts transgene expression, providing support that HUSH-mediated transcriptional repression counteracts rAAV expression in human PBMCs as well. This highlights the importance of considering host-cell regulatory mechanisms in AAV transgene design to ensure robust and durable transgene expression.

### 2.3. CD7-AAV6/9 targets T cell lines and primary human CD7^+^ lymphocytes with high specificity *in vitro*

We went on to evaluate transduction efficiency as well as off-target activity of CD7-AAV6/9 particles in T cell lines, PBMCs and a range of off-target cell types. First, intron-optimized CD7-AAV6/9 and CD7-AAV6 vectors encoding GFP were compared to their wild-type counterparts in Jurkat and SupT1 cell lines with GFP expression determined at 3 days by flow cytometry (Fig 2A, B). Notably, both cell lines express surface CD7 (Suppl Fig S1). We found that CD7-Nb engineered capsid variants exceeded transduction rates consistently compared to the wild-type variants (Fig 2A, B). Specifically, CD7-AAV6/9 achieved an over four-fold increase in GFP-expression in SupT1 cells compared to the non-engineered counterpart, while enhancement in Jurkat cells was moderate (∼1.25-fold) (Fig 2A, B).

**Figure 2:**
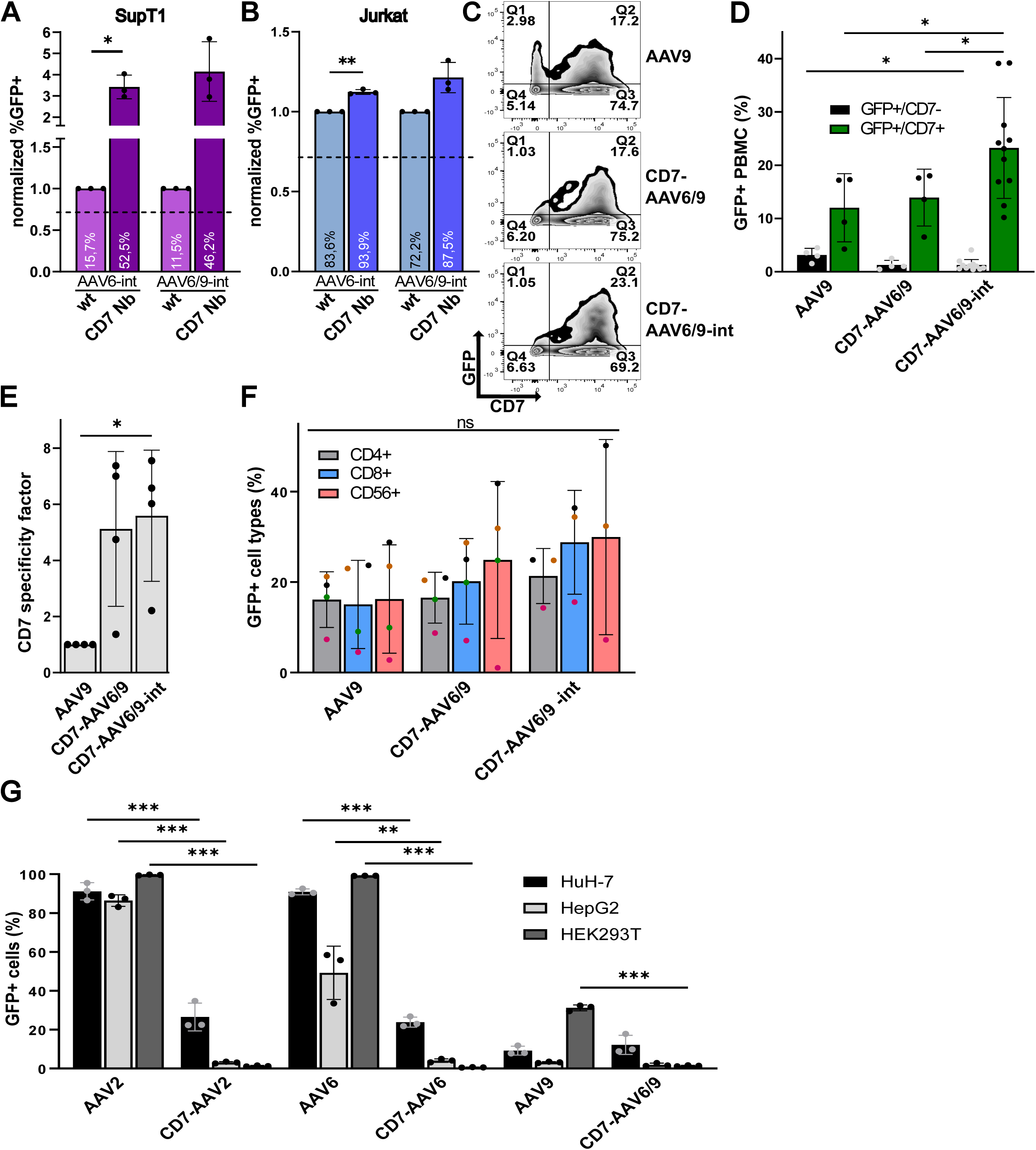
CD7-nanobody-engineered AAVs show high on-target and low off-target specificity. Transduction of SupT1 **(A)** and Jurkat **(B)** cells with wild-type (wt) and CD7-engineered AAVs of different serotypes (10,000 gc/cell). Shown are relative % GFP-positive cells at 3 dpi compared to corresponding wt AAV serotype (mean ± SD, n=3). Absolute percentage GFP-positive cells are indicated inside the bars. **C**) Representative flow cytometry plots of activated human PBMCs transduced with indicated rAAVs (100,000 gc/cell). **D)** Absolute % GFP-positivity at 3 dpi (mean ± SD, n≥4). Individual values are presented as dots. **E)** CD7-specificity factor for transduction (10,000 gc/cell) of activated PBMCs at 3 dpi (mean ± SD, n=3). **F)** Transduction of activated human PBMCs with indicated rAAVs (100,000 gc/cell) and stratification of GFP-positive cells at 3 dpi in CD4^+^ T, CD8^+^ T and NK (CD56^+^) cell types (mean ± SD, n≥3; dot colors represent biologically independent human cell samples). **G)** Transduction of hepatocyte (HuH-7, HepG2) and kidney (HEK293T) cell lines with indicated rAAVs (100.000 gc/cell). Absolute % GFP-positive cells at 3 dpi (mean ± SD, n=3). rAAV Vectors contained the following sequences: AAV6 & AAV2 = self-complementary genome, CAG promoter; AAV9 & AAV6/9 = single stranded genome, SFFV promoter. Mean ± SD, n=3. For all panels statistical analysis utilized unpaired t-test, p < 0.05 (*), p < 0.01 (**), p < 0.001 (***), and p < 0.0001 (****).

Next, we extended our analysis to primary immune cells isolated from peripheral blood (PBMCs). Intron-optimized CD7-AAV6/9 vector achieved the highest transduction efficiency, with up to 40% GFP-expressing cells compared to wild-type AAV9 and CD7-AAV6/9 without intron insertion (average 24.5%) (Mean ± SD: AAV9: 12.02 ± 6.39; CD7-AAV6/9: 13.93 ± 5.33; CD7-AAV6/9-int: 24.53 ± 8.89) (Fig 2C).

This represented an ∼2-fold increase relative to both wild-type AAV9 and CD7-AAV6/9 lacking the intron. Notably, off-target transduction of CD7-negative PBMCs was markedly reduced compared to wild-type AAV9 (Mean ± SD: AAV9 = 3.15 ± 1.24; CD7-AAV6/9: 1.27 ± 0.88; CD7-AAV6/9-int: 1.24 ± 1.10), as determined by the CD7-specificity factor (Mean ± SD: 7.04 ± 3.25 fold increase compared to wild-type AAV9) (Fig 2D).

To determine which PBMC subsets were targeted, we quantified CD4⁺ and CD8⁺ T cells as well as NK cells (CD56-positive) within the CD7/GFP double-positive population. We found that CD7-AAV6/9 transduced all CD7-positive cell populations effectively and in a comparable subset pattern to wt AAV9 (Fig 2E). We also observed that intron optimization of the rAAV transgene tended to improve targeting of T- lymphocytes and NK cells, although the data did not reach statistical significance consistent with donor-dependent variability in AAV permissivity.

Having established robust on-target activity, we next examined potential off-target effects of CD7-AAV6/9. Most wild-type AAV serotypes naturally transduce hepatocytes and renal epithelial cells due to high surface expression of proteoglycans and AAV-binding receptors in these tissues, making the liver and kidney among the most permissive organs for AAV entry [14]. We therefore exposed two hepatocyte-derived cell lines (HuH-7, HepG2) and one kidney-derived line (HEK293T) to either wild-type AAV2, 6 and 9 or their CD7-engineered capsid counterparts (Fig 2F). As expected, wild-type AAV2 and AAV6 efficiently infected all three lines. Wild-type AAV9 showed tropism only towards kidney-derived HEK293T cells. In contrast, CD7-AAV6/9-int exhibited strongly reduced transduction rates for all cell types. Interestingly, nanobody engineering of the wild-type capsids alone was sufficient to effectively abolish off-target transduction, with particularly prominent effects in HepG2 and HEK293T, showing near-complete loss in transduction (Fig 2F).

Collectively, these findings demonstrated that the intron-optimized CD7-AAV6/9 vector achieved enhanced and specific *in vitro* targeting of CD7-positive immune cells while effectively abolishing transduction of common off-target cell types.

### 2.4. Systemic targeting of human CD7-positive immune cells by intron-optimized CD7-AAV6/9 particles in humanized mice

To evaluate the *in vivo* targeting ability of intron-optimized CD7-AAV6/9 particles, we used a humanized mouse model engrafted with human PBMCs. Mice were systemically administered with three doses of rAAV encoding GFP (Fig 3A). At 3, 5, and 7 days after the last administrated dose, cells were isolated from peripheral blood, bone marrow, liver, lung, spleen and brain. Human lymphocytes were identified by flow cytometry gating (hCD45⁺/hCD3⁺), followed by assessment of CD7 expression and GFP-positivity to quantify on-target transduction (gating strategy in Suppl Fig 2). GFP-positive human cells were detected in all tissues examined, with the highest levels in the liver (>20%) and the lowest in the brain (<1%), consistent with the relative distribution of human immune cells across organs in humanized mice (Fig 3B). Analysis of the CD7 specificity of AAV transduction revealed an overall targeting towards the human CD7-positive cell compartment (Fig 3C). However, delivery across tissues was variable with a specificity factor of 4.2 in the spleen, followed by liver (3.4) and lung (2.4) (Fig 3B). At day 7, we saw a slight increase or stabilization of GFP/CD7 double-positive populations in most tissues, except for liver where percentage of GFP/CD7 double positive cells peaked at day 5 (22.05 ± 4.60%) (Fig 3C). Specificity was further established by the detection of GFP in human CD3^+^ T cells and CD56+ NK cells but not CD19^+^ B lymphocytes that do not express CD7 (Fig 3D and Suppl Fig 3).

**Figure 3:**
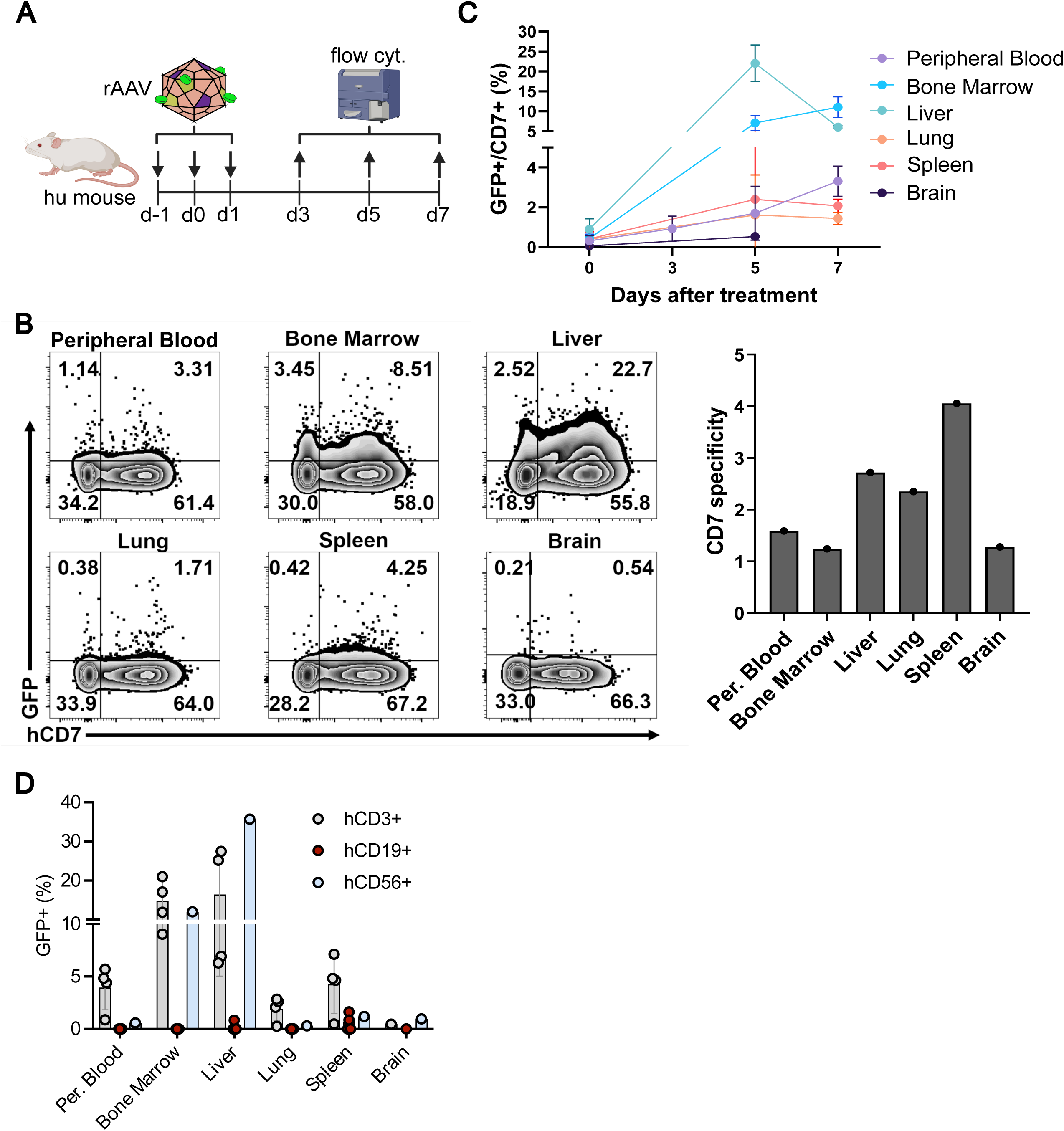
CD7-AAV6/9 vectors show efficient and specific target cell transduction in humanized mice. **A)** Study outline for CD7-AAV6/9 treatment and analysis in humanized mice. **B)** Representative flow cytometry plots depicting GFP^+^/hCD7^+^ in hCD3^+^/hCD45^+^ cells from indicated organs at 5 dpi. CD7-specificity factor are shown on the right. **C)** Percent GFP^+^/CD7^+^ cells in indicated organs at the indicated time points post treatment (mean ± SD, n≥2, except brain tissue where n = 1). **D)** Percent GFP-positive cells in indicated cell types from indicated organs at 5 days post treatment (mean ± SD, n≥2, except brain tissue where n = 1).

Taken together, CD7-AAV6/9-intron vectors mediate CD7-specific and robust transgene expression in human lymphocytes upon *in vivo* delivery in humanized mice.

## 3. Discussion

We here present a novel rAAV vector for lymphocyte-targeted gene therapy, which combines nanobody-capsid engineering and transgene cassette optimization. In detail, incorporation of a human CD7-specific nanobody into AAV6 capsid protein VP1 and complementing AAV9 VP2/3 yielded CD7-AAV6/9 particles, which are highly specific for human CD7-positive lymphocytes, including T and NK cells *in vitro* and *in vivo* with show strongly reduced off-target transduction of liver and kidney cell lines. Moreover, intron insertion into the transgene cassette boosted both transduction rates and expression levels in target cells.

This study highlights the advantages of an integrative ‘capsid-plus-genome’ vector design and opens multiple avenues for further investigation. We observed that insertion of the *HBB* intron 2 sequence boosted rAAV transgene expression in lymphocytes consistently, however it reduced the packaging capacity of rAAV vectors. Future studies focusing on investigating requirements for intron size, alternative intron sequences (endogenous or synthetic introns) and intron positioning within the transgene cassette could provide indications for additional rAAV genome optimization. Recombinant AAV transgene cassettes frequently employ promoter elements of viral origin to drive robust expression. Going forward, it could be valuable to assess human gene promoters to drive rAAV transgene expression, particularly in context of rAAVs targeted to lymphocytes that usually require cell activation for transgene expression from conventional promoters. Finally, we have previously shown that multiple sites in the capsid proteins VP1 and VP2 tolerate nanobody insertion and that even large tandem nanobody dimers can be displayed on the capsid surface [16]. This opens the possibility that combinatorial approaches incorporating different nanobodies on the same AAV capsids could further optimize target cell specificity and potentially transduction efficacy. Different receptor ligands on rAAV capsids may also boost transduction rates as multivalent ligand-receptor engagements can improve vector uptake [19].

In summary, this study supports rAAV capsid–nanobody engineering as a robust and flexible framework for generating next-generation cell-targeted vectors. Optimal vectors will combine (1) high-affinity nanobody(ies) to receptors that support specific and productive vector uptake and (2) an expression cassette designed to resist host-mediated silencing owing to its structural and functional resemblance to cellular gene context. Together, these design elements can substantially advance *in vivo* gene therapies. Notably, these efforts will rely on further improving our understanding of surface epitope composition as well as transcriptional and translational control mechanisms of designated target cells. The herein-described intron-optimized CD7-AAV6/9 vector exemplifies this integrated design and is a promising candidate for next-generation *in vivo* human immune cell gene therapy approaches that are capable of bypassing cumbersome *ex vivo* manipulation.

## 4. Materials and methods

### 4.1. Plasmids and cloning

AAV vector genomes were derived from Addgene plasmids #37825 (single-stranded) and #83279 (self-complementary). SFFV or EF1a promoters were introduced by restriction enzyme based cloning. Intron 2 of the human *HBB* gene was cloned downstream of the transgene utilizing restriction enzymes *EcoR*I/*Sal*I. Full length (850 bp) or shortened (lacking the last 60 bp, removing the branch-point site, 790 bp) intron variants were amplified from genomic DNA of primary CD4 T cells using the oligos provided in Suppl Table 1. Introns were inserted in either sense or anti-sense direction (Fig 2A).

AAV capsid variants are based on the Addgene plasmids #104963 (AAV2), #110770 (AAV6) and #112865 (AAV9), used as templates for further capsid optimization, receptor binding and mutation modifications (described below). All modifications were introduced by side-directed mutagenesis PCRs into the respective ORFs: AAV2/AAV9 - VP1 start codon mutation (ATG to AAG), AAV2/AAV6 - splice donor site mutation D9Y, AAV2 - heparan sulfate proteoglycan (HSPG) binding mutation R585A/R588A, AAV6 - HSPG binding mutation K532E and sialic acid (SIA) binding mutation V474D, AAV2 - capsid optimization modifications Y444F, T491V, Y500F, and Y730F, AAV6 - capsid optimization modifications Y705F/Y731F [16,25–32].

Human anti-CD4 nanobodies were described previously [16,33]. The human CD7 nanobody sequence was derived from the public domain ([34]; US20170226204A1) and synthesized by GenScript. Nanobodies were cloned in frame into the VP1 ORF at position T456 in VP1 (GH2/GH3 surface loop region), flanked by GSSS flexible linker (sequences provided in Suppl Table 2) [16]. All constructs were sequence verified by Sanger or long-read sequencing.

### 4.2. Cell culture

All cell lines have been previously described and were maintained at 37 °C, 5% CO₂. Adherent HEK293T (ATCC: CRL-3216), HepG2 (ATCC: HB-8065) and HuH-7 (JCRB0403) [35], cells were cultured in DMEM (Pan-Biotech) supplemented with 10% fetal calf serum (FCS; Pan Biotech), and 50 U/mL Penicillin and 50 µg/mL Streptomycin (Pan Biotech) and passaged using 0.05% trypsin-EDTA (Pan-Biotech). Suspension cell lines SupT1 (ATCC: CRL-1942) and Jurkat (RRID: CVCL_C831) were maintained in RPMI 1640 (Biozym Scientific) with 10% FCS, 50 U/mL Penicillin and 50 µg/mL Streptomycin (Pan Biotech). Peripheral blood mononuclear cells (PBMCs) were isolated from buffy coats of healthy donors (University Hospital Hamburg-Eppendorf) and cultured in RPMI 1640 with 10% FCS, 50 U/mL Penicillin and 50 µg/mL Streptomycin (Pan Biotech), 200 U/mL recombinant human IL-2 (Sigma-Aldrich). PBMCs were activated with ImmunoCult hCD3/CD28 T cell activation reagent (StemCell Technologies) three days before transduction according to manufacturer’s instructions. Work including primary human cells were conducted in accordance with ethical guidelines and approved by the local ethics committee (Ärztekammer Hamburg; PV4666 and WF010/2011).

### 4.3. AAV production

AAV production using polyethylenimine (PEI; Sigma-Aldrich) transfection in HEK293T cells (ATCC: CRL-3216) as described before with minor modifications depending on the serotype produced [16]. Briefly, 4 × 10⁶ cells per 10 cm dish were seeded one day prior transfection. Co-transfection was conducted in equimolar ratios of the helper plasmid pAdDeltaF6 (Addgene #112867), vector genome (single-stranded (Addgene #37825) or self-complementary (Addgene #83279)) and Rep/Cap plasmids encoding VP1-Nanobody and VP2/VP3 proteins, respectively (4:1 PEI:DNA ratio, total of 10.5 μg plasmid DNA per dish). For AAV9 and hybrid AAV6/9 virus was harvested from supernatant: medium was collected 48 hours post transfection and stored at 4 °C. Six mL of fresh complete medium was added and supernatant again collected 96 hours post transfection and pooled. Supernatant was cleared by centrifugation (1699 g, 15 min, 4 °C) and filtered through a 0.22 µm maze. Particles were precipitated using PEG (final 10% PEG8000 ,1M NaCl) for 48 h at 4 °C. Precipitated virus was pelleted by centrifugation (1699 g, 30 min, 4 °C), resuspended in 800 µl PBS and stored at −70 °C. AAV2 and 6 were isolated from cell lysates: transfected cells were harvested 96 hours post transfection by scraping, pelleted by centrifugation (15 min, 2,800 g) and resuspended in 800 μL lysis buffer (150 mM NaCl, 50 mM Tris-HCl pH 8.0 in water). Four freeze/thaw cycles (−70 °C, 20 min; 37 °C, 5 min) released AAV particles from the producer cells. Subsequent benzonase treatment (50 U, 37 °C, 1 h; Sigma-Aldrich) reduced contaminating nucleic acid content. Cell debris was removed by centrifugation (20,000 g, 4 °C, 15 min), yielding a crude lysate, stored at −70 °C. For further purification AAV vectors were purified by iodixanol gradient ultracentrifugation [16,36]. Briefly, 10 mL of AAV stock was layered onto an iodixanol step-gradient and ultracentrifuged (350,000 × g, 90 min, 4 °C). The 40% iodixanol fraction containing purified AAV was collected, concentrated and buffer exchanged (PBS supplemented with 0.001% Pluronic F-68 (Gibco)) using centrifugal filters (Amicon Ultra-15, 100 kDa MWCO; Sigma-Aldrich). Purified vector preparations were aliquoted and stored at −70 °C.

### 4.4. Vector Genome Quantification

Vector titers were determined by qPCR [16]. Briefly, 1 μL AAV vector stock was treated with DNaseI to eliminate residual plasmid DNA (10 U DNaseI (NEB), 100 µL reaction, 37 °C,16 h), followed by DNaseI inactivation (75 °C, 30 min) and Proteinase K digest to release viral genomes (40 ug Proteinase K (QIAGEN), 55 °C,2 h). Proteinase K was inactivated (95 °C, 30 min. One μL of this reaction was subjected to qPCR (PowerUp SYBR Green Master Mix,Applied Biosystems 7500 real-time PCR system) using plasmid standards rows for absolute copy number determination. Primer sequences are provided in Suppl Table 1.

AAV genome delivery was quantified by determining vector genome copy numbers in genomic DNA (gDNA) isolated from infected T cells and normalized to human β-globin (HBG) gene copies in the same samples. Infected and non-infected cells were FACS sorted and gDNA isolated using QIAamp DNA Blood Mini Kit (Qiagen). Vector genome copy numbers were determined by qPCR. HBG copy numbers were quantified by TaqMan qPCR. AAV vector genome copy numbers were then normalized to HBG copy numbers, yielding viral genome copies per cell. Primer sequences are provided in Suppl Table 1.

### 4.5. AAV transduction and flow cytometry

Cells were infected with indicated AAV variants at defined genome copies per cell (gc/cell), for each experiment. Activated PBMCs were seeded at 4 × 10⁴ cells per well in 96U-well plates in 30 µL culture medium before adding the AAV stocks supplemented with 50 µL RPMI/IL-2 5 hours post transduction. Three days post transduction (dpi), PBMCs were washed with PBS and stained for surface markers in PBS containing 2% FCS and 1 mM EDTA (1 h, 4 °C,) using PE-Cy7–conjugated anti-CD4 (RPA-T4; eBioscience), APC–conjugated anti-CD7 (M-T701; BD Biosciences), BV421–conjugated anti-CD8 (RPA-T8; BioLegend), and BV605–conjugated anti-CD56 (NCAM 16.2; BD Biosciences). After two PBS washes cells were analyzed for CD4, CD7, CD8, CD56, and GFP expression by flow cytometry. For AAV transductions in cell lines, 4 × 10⁴ HepG2, HuH-7, SupT1 and Jurkat cells were seeded in 96-well plates and exposed to AAV at the indicated gc/cell. Prior to measurement, cells were washed twice with PBS and fixed with 0.5% paraformaldehyde in PBS. Data acquired on Canto II or LSR Fortessa (BD Biosciences).

### 4.6. Studies in humanized mice

NSG-Tg(Hu-IL15) or NSG-15 mice (Strain #:030890 purchased from the Jackson Laboratory, Bar Harbor, ME, USA) were bred and maintained under pathogen-free conditions at the Yale Animal Resource Center. All animal studies were approved by the Yale Institutional Animal Care and Use Committees (IACUC). NSG-15 were engrafted with PBMC as described previously [37,38]. Briefly, 3.5 × 10^6^ PBMCs, purified by Ficoll density gradient centrifugation of healthy donor blood buffy coats (obtained from the New York Blood Bank) were injected intraperitoneally in a 200 µL volume into 6- to 8-week-old NSG-15 mice (both male and female), using a 1-cm^3^ syringe and a 25-gauge needle. 2-3 weeks after transplantation, PBMCs from 100 µl blood obtained by Ficoll density gradient centrifugation were stained with fluorochrome-conjugated anti-human CD45-APC (HI30; BD Biosciences), CD3-Alexa Fluor 700 (UCHT1; BioLegend), CD4-PerCP (OKT4; BioLegend), CD8-V500 (SK1; BD Biosciences), and CD19-BV421 (SJ25C1; BioLegend) antibodies, and analyzed by flow cytometry to confirm human cell engraftment. Humanized mice were intravenously administered (through the retro-orbital route) CD7-AAV-6/9-int-GFP at a dose of 5×10^10^ vg/mouse in 100 µl volume on three consecutive days. GFP expression was analyzed in peripheral blood cell subsets on days 3, 5, and 7 after treatment by flow cytometry as described above. GFP expression was also analyzed in cells from tissues obtained on day 5 post treatment. Tissues were minced with scissors and digested with DNaseI (10 IU/mL) and CollagenaseD (1 mg/mL) at 37 °C for 30 min with gentle shaking. The digested samples were washed, and cells were isolated using 33% Percoll in RPMI1640. Red blood cells were lysed with ACK lysis buffer and the remaining cells were stained with fluorochrome-conjugated anti-human CD45-APC (HI30; BD Biosciences), CD3-Alexa Fluor 700 (UCHT1; BioLegend), CD4-PerCP (OKT4; BioLegend), CD8-V500 (SK1; BD Biosciences), and CD7-PE (124-1D1; Invitrogen) antibodies and analyzed by flow cytometry to assess GFP expression.

### 4.7. Electron microscopy and immuno-gold staining

Iodixanol-purified CD7-AAV6/9 preparations were adsorbed onto glow-discharged, carbon-coated grids (10 min). Grids were blocked (5 min) with blocking solution for protein A/G gold conjugates (AURION) and incubated with primary antibody (10 μg/mL polyclonal goat anti-alpaca IgG; Jackson ImmunoResearch; 30 min), followed by 5 min wash. Secondary labeling was performed with rabbit anti-goat IgG conjugated to 10 nm gold particles (1:10 dilution; Sigma-Aldrich; 15 min). After additional washes with ddH₂O, grids were negatively stained with 1% uranyl acetate and air-dried. Transmission electron microscopy was performed using a FEI Tecnai G20 operated at 80 kV, equipped with an Olympus Veleta CCD camera. Representative micrographs were cropped for clarity, and scale bars were included.

### 4.8. Statistical analysis and software

Flow cytometry data were analyzed using FlowJo v10.8.1 (Tree Star) and processed in GraphPad Prism v10.2.2 (GraphPad Software). Data are presented as means ± SD; the number of independent experiments is indicated in the figure legends. Statistical significance was assessed using unpaired *t*-tests in GraphPad Prism with *p*-values denoted as: *p* < 0.05 (\**), p < 0.01 (**), p < 0.001 (\*\*\**), and *p* < 0.0001 (****). The CD7 specificity factor was calculated as: (%GFP+ cells in CD7+ population) / (%GFP+ cells in CD7- population) [16].

### 4.9. Protein structure modeling

Structure prediction modeling using AlphaFold was used to compare AAV6 VP1 with and without CD7 nanobody insertion [39]. Input sequences for the AF3 server of AAV6 VP1 with nanobody (886 aa) and without nanobody (736 aa) are provided in Suppl Table 2. 3D modelled structures were superimposed using Matchmaker structure analysis tool in UCSF ChimeraX [40–42]. Briefly, MatchMaker first generates secondary structure based pairwise sequence alignment followed by superimposing the 3D coordinates of the respective Cα atoms. Alignment was performed with default parameters.

## Acknowledgments

This research is supported by BMFTR grant number 01KI2105, TheraImmun project funded by EFRE and IFB, NIAID award numbers R01AI181053 (awarded to U.C.L and P.K.), R01AI145164 (awarded to P.K.) and UM1AI164559, co-funded by NHLBI, NIDA, NIMH, NINDS, and NIDDK. The LIV is supported by the Free and Hanseatic City of Hamburg and the Federal Ministry of Health. The authors thank Carola Schneider and Rudolph Reimer (LIV) for providing EM images as well as Arne Düsedau and Jana Hennesen (LIV Flow Cytometry Core facility).

## Author contributions

U.C.L. and M.V.H. conceived the study; H.J. conducted the *in vitro* experiments; H.K., Y.S. conducted the *in vivo* experiments; U.E.S. performed AlphaFold modeling; M.V.H., N.S.Q., D.F., N.B. supported experimental work; H.J., M.V.H., P.K. and U.C.L. designed the study and interpreted the data; H.J., M.V.H., U.C.L., P.K. wrote the manuscript; H.K., Y.S., D.F., N.B. revised the manuscript; U.C.L., P.K. provided funding.

## Declaration of interest

The authors declare no competing interests.

## Notes

### Competing Interest Statement

The authors have declared no competing interest.

## References

[1] S. Russell, J. Bennett, J.A. Wellman, D.C. Chung, Z.F. Yu, A. Tillman, J. Wittes, J. Pappas, O. Elci, S. McCague, D. Cross, K.A. Marshall, J. Walshire, T.L. Kehoe, H. Reichert, M. Davis, L. Raffini, L.A. George, F.P. Hudson, L. Dingfield, X. Zhu, J.A. Haller, E.H. Sohn, V.B. Mahajan, W. Pfeifer, M. Weckmann, C. Johnson, D. Gewaily, A. Drack, E. Stone, K. Wachtel, F. Simonelli, B.P. Leroy, J.F. Wright, K.A. High, A.M. Maguire, Efficacy and safety of voretigene neparvovec (AAV2-hRPE65v2) in patients with RPE65-mediated inherited retinal dystrophy: a randomised, controlled, open-label, phase 3 trial, Lancet (London, England) 390 (2017) 849–860. 10.1016/S0140-6736(17)31868-8.

[2] D. Gaudet, J. Méthot, S. Déry, D. Brisson, C. Essiembre, G. Tremblay, K. Tremblay, J. De Wal, J. Twisk, N. Van Den Bulk, V. Sier-Ferreira, S. Van Deventer, Efficacy and long-term safety of alipogene tiparvovec (AAV1-LPLS447X) gene therapy for lipoprotein lipase deficiency: an open-label trial, Gene Ther. 2013 204 20 (2012) 361–369. 10.1038/gt.2012.43.

[3] J.W. Day, R.S. Finkel, C.A. Chiriboga, A.M. Connolly, T.O. Crawford, B.T. Darras, S.T. Iannaccone, N.L. Kuntz, L.D.M. Peña, P.B. Shieh, E.C. Smith, J.M. Kwon, C.M. Zaidman, M. Schultz, D.E. Feltner, S. Tauscher-Wisniewski, H. Ouyang, D.H. Chand, D.M. Sproule, T.A. Macek, J.R. Mendell, Onasemnogene abeparvovec gene therapy for symptomatic infantile-onset spinal muscular atrophy in patients with two copies of SMN2 (STR1VE): an open-label, single-arm, multicentre, phase 3 trial, Lancet Neurol. 20 (2021) 284–293. 10.1016/S1474-4422(21)00001-6.

[4] J.H. Wang, D.J. Gessler, W. Zhan, T.L. Gallagher, G. Gao, Adeno-associated virus as a delivery vector for gene therapy of human diseases, Signal Transduct. Target. Ther. 2024 91 9 (2024) 1–33. 10.1038/s41392-024-01780-w.

[5] A.C. Nathwani, U.M. Reiss, E.G.D. Tuddenham, C. Rosales, P. Chowdary, J. McIntosh, M. Della Peruta, E. Lheriteau, N. Patel, D. Raj, A. Riddell, J. Pie, S. Rangarajan, D. Bevan, M. Recht, Y.-M. Shen, K.G. Halka, E. Basner-Tschakarjan, F. Mingozzi, K.A. High, J. Allay, M.A. Kay, C.Y.C. Ng, J. Zhou, M. Cancio, C.L. Morton, J.T. Gray, D. Srivastava, A.W. Nienhuis, A.M. Davidoff, Long-Term Safety and Efficacy of Factor IX Gene Therapy in Hemophilia B, N. Engl. J. Med. 371 (2014) 1994–2004. 10.1056/NEJMOA1407309;ISSUE:ISSUE:DOI.

[6] H.K.E. Au, M. Isalan, M. Mielcarek, Gene Therapy Advances: A Meta-Analysis of AAV Usage in Clinical Settings, Front. Med. 0 (2022) 2746. 10.3389/FMED.2021.809118.

[7] M.F. Naso, B. Tomkowicz, W.L. Perry, W.R. Strohl, Adeno-Associated Virus (AAV) as a Vector for Gene Therapy, BioDrugs 31 (2017) 317–334. 10.1007/s40259-017-0234-5.

[8] B.C. Schnepp, J.D. Chulay, G.J. Ye, T.R. Flotte, B.C. Trapnell, P.R. Johnson, Recombinant Adeno-Associated Virus Vector Genomes Take the Form of Long-Lived, Transcriptionally Competent Episomes in Human Muscle, Https://Home.Liebertpub.Com/Hum 27 (2015) 32–42. 10.1089/HUM.2015.136.

[9] D. Wang, P.W.L. Tai, G. Gao, Adeno-associated virus vector as a platform for gene therapy delivery, Nat. Rev. Drug Discov. 18 (2019) 358–378. 10.1038/s41573-019-0012-9.

[10] A. Westhaus, M. Cabanes-Creus, A. Rybicki, G. Baltazar, R.G. Navarro, E. Zhu, M. Drouyer, M. Knight, R.F. Albu, B.H. Ng, P. Kalajdzic, M. Kwiatek, K. Hsu, G. Santilli, W. Gold, B. Kramer, A. Gonzalez-Cordero, A.J. Thrasher, I.E. Alexander, L. Lisowski, High-Throughput In Vitro , Ex Vivo, and In Vivo Screen of Adeno-Associated Virus Vectors Based on Physical and Functional Transduction, Hum. Gene Ther. 31 (2020) 575–589. 10.1089/hum.2019.264.

[11] M. Penaud-Budloo, C. Le Guiner, A. Nowrouzi, A. Toromanoff, Y. Chérel, P. Chenuaud, M. Schmidt, C. von Kalle, F. Rolling, P. Moullier, R.O. Snyder, Adeno-Associated Virus Vector Genomes Persist as Episomal Chromatin in Primate Muscle, J. Virol. 82 (2008) 7875–7885. 10.1128/jvi.00649-08.

[12] C. Hagedorn, M. Schnödt-Fuchs, P. Boehme, H. Abdelrazik, H.J. Lipps, H. Büning, S/MAR Element Facilitates Episomal Long-Term Persistence of Adeno-Associated Virus Vector Genomes in Proliferating Cells, in: Hum. Gene Ther., Hum Gene Ther, 2017: pp. 1169–1179. 10.1089/hum.2017.025.

[13] B.L. Ellis, M.L. Hirsch, J.C. Barker, J.P. Connelly, R.J. Steininger, M.H. Porteus, A survey of ex vivo/in vitro transduction efficiency of mammalian primary cells and cell lines with Nine natural adeno-associated virus (AAV1-9) and one engineered adeno-associated virus serotype, Virol. J. 10 (2013) 74. 10.1186/1743-422X-10-74.

[14] A. Srivastava, In vivo tissue-tropism of adeno-associated viral vectors, Curr. Opin. Virol. 21 (2016) 75–80. 10.1016/j.coviro.2016.08.003.

[15] D. Duan, Lethal immunotoxicity in high-dose systemic AAV therapy, Mol. Ther. 31 (2023) 3123–3126. 10.1016/j.ymthe.2023.10.015.

[16] M. V. Hamann, N. Beschorner, X.-K. Vu, I. Hauber, U.C. Lange, B. Traenkle, P.D. Kaiser, D. Foth, C. Schneider, H. Büning, U. Rothbauer, J. Hauber, Improved targeting of human CD4+ T cells by nanobody-modified AAV2 gene therapy vectors, PLoS One 16 (2021) e0261269. 10.1371/journal.pone.0261269.

[17] D.W. Russell, M.A. Kay, Adeno-Associated Virus Vectors and Hematology, Blood 94 (1999) 864. 10.1182/blood.v94.3.864.415k34_864_874.

[18] M.B. Demircan, L.J. Zinser, A. Michels, M. Guaza-Lasheras, F. John, J.M. Gorol, S.A. Theuerkauf, D.M. Günther, D. Grimm, F.R. Greten, P. Chlanda, F.B. Thalheimer, C.J. Buchholz, T-cell specific in vivo gene delivery with DART-AAVs targeted to CD8, Mol. Ther. 32 (2024). 10.1016/j.ymthe.2024.08.002.

[19] S.A. Theuerkauf, E. Herrera-Carrillo, F. John, L.J. Zinser, M.A. Molina, V. Riechert, F.B. Thalheimer, K. Börner, D. Grimm, P. Chlanda, B. Berkhout, C.J. Buchholz, AAV vectors displaying bispecific DARPins enable dual-control targeted gene delivery, Biomaterials 303 (2023). 10.1016/j.biomaterials.2023.122399.

[20] O. Olarewaju, F. Held, P. Curtis, C.H. Kenny, U. Maier, T. Panavas, F. du Plessis, αFAP-specific nanobodies mediate a highly precise retargeting of modified AAV2 capsids thereby enabling specific transduction of tumor tissues, Mol. Ther. Methods Clin. Dev. 32 (2024). 10.1016/j.omtm.2024.101378.

[21] M.D. Hoffmann, J.P. Gallant, A.M. LeBeau, D. Schmidt, Unlocking precision gene therapy: harnessing AAV tropism with nanobody swapping at capsid hotspots, NAR Mol. Med. 1 (2024). 10.1093/narmme/ugae008.

[22] A.M. Eichhoff, K. Börner, B. Albrecht, W. Schäfer, N. Baum, F. Haag, J. Körbelin, M. Trepel, I. Braren, D. Grimm, S. Adriouch, F. Koch-Nolte, Nanobody-Enhanced Targeting of AAV Gene Therapy Vectors, Mol. Ther. - Methods Clin. Dev. 15 (2019) 211–220. 10.1016/j.omtm.2019.09.003.

[23] D. Grimm, J.S. Lee, L. Wang, T. Desai, B. Akache, T.A. Storm, M.A. Kay, In Vitro and In Vivo Gene Therapy Vector Evolution via Multispecies Interbreeding and Retargeting of Adeno-Associated Viruses, J. Virol. 82 (2008) 5887–5911. 10.1128/jvi.00254-08.

[24] R.C. Münch, H. Janicki, I. Völker, A. Rasbach, M. Hallek, H. Büning, C.J. Buchholz, Displaying high-affinity ligands on adeno-associated viral vectors enables tumor cell-specific and safe gene transfer, Mol. Ther. 21 (2013) 109–118. 10.1038/mt.2012.186.

[25] H. Büning, A. Srivastava, Capsid Modifications for Targeting and Improving the Efficacy of AAV Vectors, Mol. Ther. - Methods Clin. Dev. 12 (2019) 248–265. 10.1016/J.OMTM.2019.01.008.

[26] A. Kern, K. Schmidt, C. Leder, O.J. Müller, C.E. Wobus, K. Bettinger, C.W. Von der Lieth, J.A. King, J.A. Kleinschmidt, Identification of a Heparin-Binding Motif on Adeno-Associated Virus Type 2 Capsids, J. Virol. 77 (2003) 11072–11081. 10.1128/jvi.77.20.11072-11081.2003.

[27] S.R. Opie, K.H. Warrington, M. Agbandje-McKenna, S. Zolotukhin, N. Muzyczka, Identification of Amino Acid Residues in the Capsid Proteins of Adeno-Associated Virus Type 2 That Contribute to Heparan Sulfate Proteoglycan Binding, J. Virol. 77 (2003) 6995–7006. 10.1128/jvi.77.12.6995-7006.2003.

[28] J. O’Donnell, K.A. Taylor, M.S. Chapman, Adeno-associated virus-2 and its primary cellular receptor--Cryo-EM structure of a heparin complex, Virology 385 (2009) 434–443. 10.1016/J.VIROL.2008.11.037.

[29] R. Sayroo, D. Nolasco, Z. Yin, Y. Colon-Cortes, M. Pandya, C. Ling, G. Aslanidi, Development of novel AAV serotype 6 based vectors with selective tropism for human cancer cells, Gene Ther. 23 (2016) 18–25. 10.1038/gt.2015.89.

[30] C. Ling, K. Bhukhai, Z. Yin, M. Tan, M.C. Yoder, P. Leboulch, E. Payen, A. Srivastava, High-Efficiency Transduction of Primary Human Hematopoietic Stem/Progenitor Cells by AAV6 Vectors: Strategies for Overcoming Donor-Variation and Implications in Genome Editing, Sci. Rep. 6 (2016) 1–8. 10.1038/srep35495.

[31] S. Muralidhar, S.P. Becerra, J.A. Rose, Site-directed mutagenesis of adeno-associated virus type 2 structural protein initiation codons: effects on regulation of synthesis and biological activity., J. Virol. 68 (1994) 170–6. http://www.ncbi.nlm.nih.gov/pubmed/8254726 (accessed November 20, 2019).

[32] J. Judd, F. Wei, P.Q. Nguyen, L.J. Tartaglia, M. Agbandje-Mckenna, J.J. Silberg, J. Suh, Random insertion of mcherry into VP3 domain of adeno-associated virus yields fluorescent capsids with no loss of infectivity, Mol. Ther. - Nucleic Acids 1 (2012) e54. 10.1038/mtna.2012.46.

[33] B. Traenkle, P.D. Kaiser, S. Pezzana, J. Richardson, M. Gramlich, T.R. Wagner, D. Seyfried, M. Weldle, S. Holz, Y. Parfyonova, S. Nueske, A.M. Scholz, A. Zeck, M. Jakobi, N. Schneiderhan-Marra, M. Schaller, A. Maurer, C. Gouttefangeas, M. Kneilling, B.J. Pichler, D. Sonanini, U. Rothbauer, Single-Domain Antibodies for Targeting, Detection, and In Vivo Imaging of Human CD4+ Cells, Front. Immunol. 12 (2021). 10.3389/fimmu.2021.799910.

[34] L. Yang, J. Tang, CD7-Nb patent US20170226204A1, 2014.

[35] H. Nakabayashi, K. Taketa, K. Miyano, T. Yamane, J. Sato, Growth of human hepatoma cells lines with differentiated functions in chemically defined medium., Cancer Res. 42 (1982) 3858–3863.

[36] J. Fakhiri, M. Nickl, D. Grimm, Rapid and Simple Screening of CRISPR Guide RNAs (gRNAs) in Cultured Cells Using Adeno-Associated Viral (AAV) Vectors, in: Methods Mol. Biol., Humana Press Inc., 2019: pp. 111–126. 10.1007/978-1-4939-9170-9_8.

[37] J. Richard, G. Sannier, L. Zhu, J. Prévost, L. Marchitto, M. Benlarbi, G. Beaudoin-Bussières, H. Kim, Y. Sun, D. Chatterjee, H. Medjahed, C. Bourassa, G.-G. Delgado, M. Dubé, F. Kirchhoff, B.H. Hahn, P. Kumar, D.E. Kaufmann, A. Finzi, CD4 downregulation precedes Env expression and protects HIV-1-infected cells from ADCC mediated by non-neutralizing antibodies., MBio 15 (2024) e0182724. 10.1128/mbio.01827-24.

[38] J.K. Rajashekar, J. Richard, J. Beloor, J. Prévost, S.P. Anand, G. Beaudoin-Bussières, L. Shan, D. Herndler-Brandstetter, G. Gendron-Lepage, H. Medjahed, C. Bourassa, F. Gaudette, I. Ullah, K. Symmes, A. Peric, E. Lindemuth, F. Bibollet-Ruche, J. Park, H.-C. Chen, D.E. Kaufmann, B.H. Hahn, J. Sodroski, M. Pazgier, R.A. Flavell, A.B. Smith, A. Finzi, P. Kumar, Modulating HIV-1 envelope glycoprotein conformation to decrease the HIV-1 reservoir., Cell Host Microbe 29 (2021) 904–916.e6. 10.1016/j.chom.2021.04.014.

[39] J. Abramson, J. Adler, J. Dunger, R. Evans, T. Green, A. Pritzel, O. Ronneberger, L. Willmore, A.J. Ballard, J. Bambrick, S.W. Bodenstein, D.A. Evans, C.C. Hung, M. O’Neill, D. Reiman, K. Tunyasuvunakool, Z. Wu, A. Žemgulytė, E. Arvaniti, C. Beattie, O. Bertolli, A. Bridgland, A. Cherepanov, M. Congreve, A.I. Cowen-Rivers, A. Cowie, M. Figurnov, F.B. Fuchs, H. Gladman, R. Jain, Y.A. Khan, C.M.R. Low, K. Perlin, A. Potapenko, P. Savy, S. Singh, A. Stecula, A. Thillaisundaram, C. Tong, S. Yakneen, E.D. Zhong, M. Zielinski, A. Žídek, V. Bapst, P. Kohli, M. Jaderberg, D. Hassabis, J.M. Jumper, Accurate structure prediction of biomolecular interactions with AlphaFold 3, Nat. 2024 6308016 630 (2024) 493–500. 10.1038/s41586-024-07487-w.

[40] E.C. Meng, T.D. Goddard, E.F. Pettersen, G.S. Couch, Z.J. Pearson, J.H. Morris, T.E. Ferrin, UCSF ChimeraX: Tools for structure building and analysis, Protein Sci. 32 (2023) e4792. 10.1002/PRO.4792;WGROUP:STRING:PUBLICATION.

[41] E.F. Pettersen, T.D. Goddard, C.C. Huang, E.C. Meng, G.S. Couch, T.I. Croll, J.H. Morris, T.E. Ferrin, UCSF ChimeraX: Structure visualization for researchers, educators, and developers, Protein Sci. 30 (2021) 70–82. 10.1002/PRO.3943.

[42] T.D. Goddard, C.C. Huang, E.C. Meng, E.F. Pettersen, G.S. Couch, J.H. Morris, T.E. Ferrin, UCSF ChimeraX: Meeting modern challenges in visualization and analysis, Protein Sci. 27 (2018) 14–25. 10.1002/PRO.3235.

[43] M. Seczynska, P.J. Lehner, The sound of silence: mechanisms and implications of HUSH complex function, Trends Genet. 39 (2023) 251–267. 10.1016/J.TIG.2022.12.005.

[44] M. Seczynska, S. Bloor, S.M. Cuesta, P.J. Lehner, Genome surveillance by HUSH-mediated silencing of intronless mobile elements, Nature 601 (2022) 440–445. 10.1038/s41586-021-04228-1.

