## Supplementary figures and images for "Development of a recombinant adeno-associated virus vector for human T lymphocyte- and natural killer cell-targeted gene therapy"

### Suppl Figure 1-3

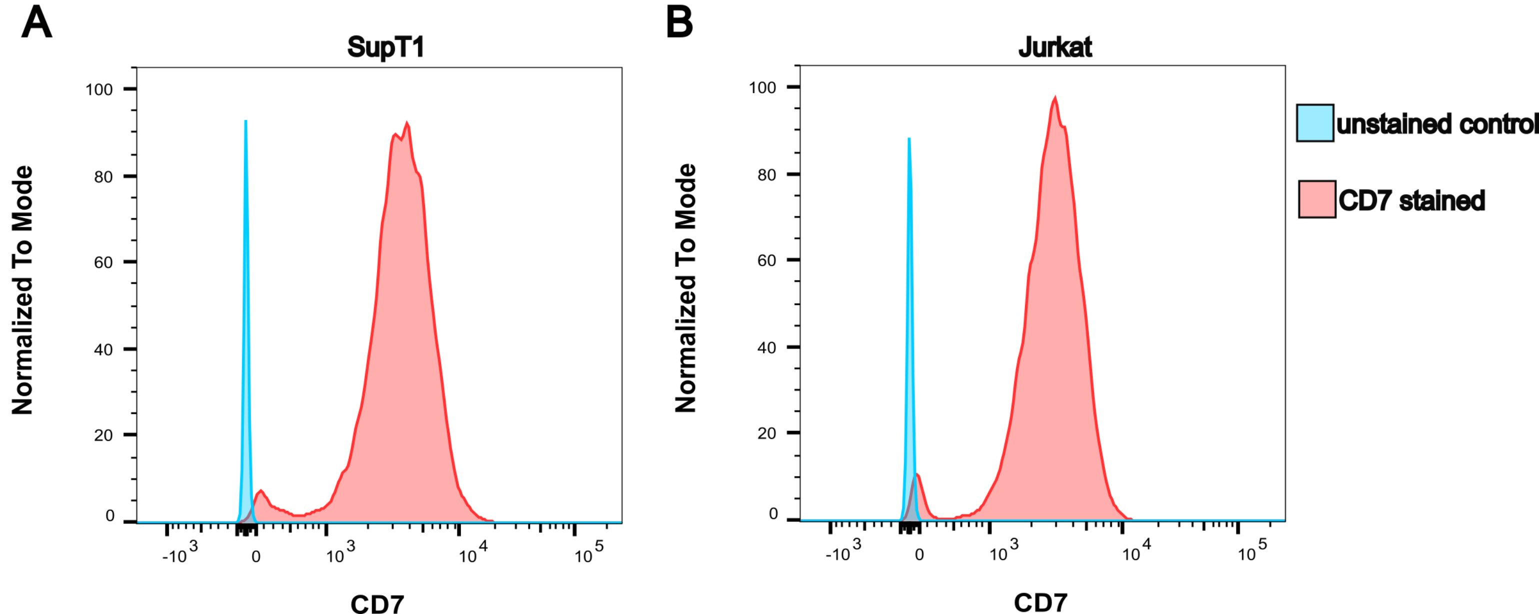

#### Figure S2

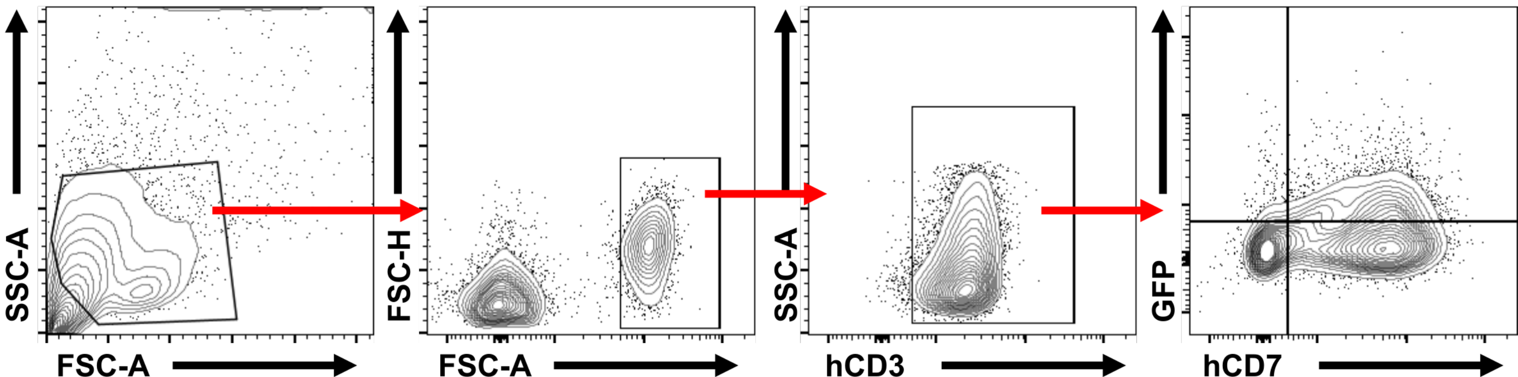

#### Figure S3

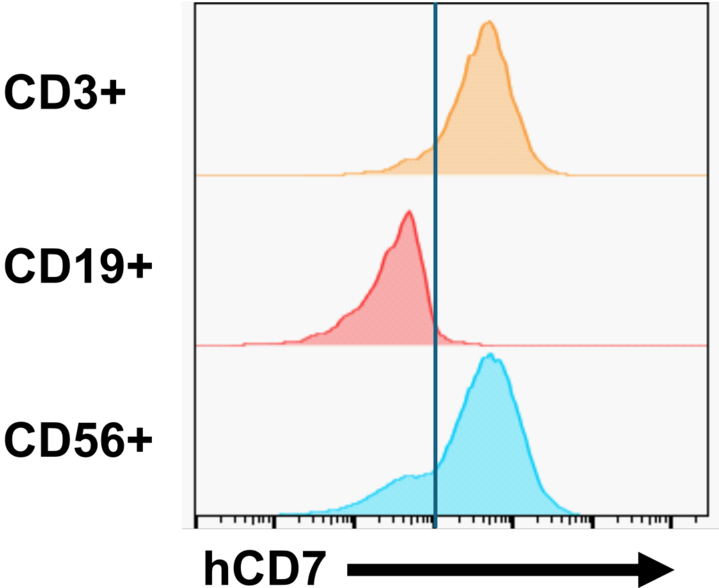
